# Robust design capture-recapture analysis of abundance and demographic parameters of Indian River Lagoon common bottlenose dolphins (*Tursiops truncatus truncatus*)

**DOI:** 10.1101/2020.01.30.926683

**Authors:** Wendy Noke Durden, Eric D. Stolen, Lydia Moreland, Elisabeth Howells, Teresa Jablonski, Anne Sleeman, Matthew Denny, George Biedenbach, Marilyn Mazzoil

## Abstract

Accurate estimates of abundance are critical to species management and conservation. Common bottlenose dolphins (*Tursiops truncatus truncatus*) inhabiting the Indian River Lagoon (IRL) estuarine system along the east coast of Florida are impacted by anthropogenic activities and have had multiple unexplained mortality events, necessitating precise estimates of demographic and abundance parameters to implement management strategies. Mark-recapture methodology following a Robust Design survey was used to estimate abundance, adult survival, and temporary emigration for the IRL estuarine system stock of bottlenose dolphins. Models included a parameter (time since first capture) to assess evidence for transient individuals. Boat-based photo-identification surveys (*n* = 135) were conducted along predetermined contour and transect lines throughout the entire IRL (2016-2017). The best fitting model included the “transient” parameter to survival, allowed survival to vary by primary period, detection to vary by secondary session, and did not allow temporary emigration. Dolphin abundance ranged from 981 (95% CI: 882-1,090) in winter to 1,078 (95% CI: 968-1,201) in summer with a mean of 1,032 (95% CI: 969 -1,098). Model averaged seasonal survival rate for marked residents ranged from 0.85-1.00. Capture probability ranged from 0.20 to 0.42 during secondary sessions and transient rate from 0.06 to 0.07. This study represents the first Robust design mark-recapture survey effort to estimate abundance for IRL dolphins and provides parameter estimates to optimize sampling design of future studies. Transients included individuals with home ranges extending north of the IRL requiring further assessment of stock delineation. Results were remarkably similar to prior abundance estimates resulting from line-transect aerial surveys and were consistent with a stable population. Data will enable managers to evaluate the impact of fisheries-related takes as well as enable future comparisons of demographic parameters for a dolphin population that continues to sustain large scale mortality events and anthropogenic impacts.

## Introduction

Anthropogenic impacts continue to threaten both coastal and estuarine cetacean populations [1–9], rendering the accurate evaluation of abundance and demographic parameters critical to population management. Various methods have been employed to estimate the abundance of cetacean species, most of which have relied on line-transect or mark-recapture methodology [10–15]. While line-transect surveys (aerial and vessel based) are likely the most utilized methods for coastal species, they are incapable of distinguishing between resident and transient animals. Identifying resident animals within a population is critical, as those animals are most vulnerable to the impacts of repeated anthropogenic activities as well as ecological declines in the region.

Common bottlenose dolphins (*Tursiops truncatus truncatus*) inhabiting the Indian River Lagoon (IRL) estuarine system along the east coast of Florida between Ponce Inlet and Jupiter Inlet have been studied for decades and are considered long-term residents comprising the IRL estuarine system dolphin stock [16–18]. However, more recent studies support the occurrence of transients as well as ranging patterns that extend beyond the northern boundary of the IRL [19–22]. The expansive range (∼250 km) of this dolphin stock has made vessel-based abundance estimation difficult and prior studies have employed line-transect aerial surveys [19, 23, 24]. While aerial surveys provide accurate and unbiased estimates of abundance and distribution, they do not provide estimates of immigration/emigration or survival which are needed for stock management. Mark-recapture techniques are widely used to estimate abundance in animals that can be marked or otherwise identified and can provide these important parameter estimates [25].

Many cetacean species, including bottlenose dolphins, can be individually identified by natural occurring markings on the trailing edge of the dorsal fin [26, 27, 28]. Photo-identification, or the identification of individuals based on these unique markings, is widely used to study cetaceans [28]. Photo-identification is commonly combined with mark-recapture methods where marked individuals are “captured” (first identified) and subsequently “recaptured” (resighted) during survey efforts to estimate abundance and demographic parameters including survival and temporary emigration [29–34].

Accurate estimates of abundance and demographic parameters are essential to the management and conservation of the IRL dolphin stock and have become increasingly important as IRL dolphins have experienced multiple Unusual Mortality Events (UMEs) (2001, 2008, 2013; 2013-2015) [35]. During the largest mortality event (2013 UME), a minimum of 77 dolphin mortalities occurred. Based on the mean abundance estimate prior to the event (1,032 dolphins) [19], mortalities represented ∼7.5% of population. Concurrent with this event, the Mid Atlantic UME (2013-2015) coincided and further impacted IRL dolphins [35]. Reoccurring mortality events could indicate serious ecological pressures that may lead to the decline of this stock. Indian River Lagoon dolphins are listed as a strategic stock since anthropogenic mortality likely exceeds Potential Biological Removal (PBR); the maximum number of mortalities (excluding natural mortalities) that can be removed annually while still allowing the stock to reach or maintain an optimal sustainable population level. In recent years, the IRL has undergone several large scale ecosystem changes, most notably phytoplankton blooms that yielded catastrophic seagrass loss [36]. Seagrass meadows have been found to provide critical habitat to prey consumed by estuarine dolphins [37], therefore these significant ecological changes are likely to jeopardize the health of the already vulnerable IRL dolphin stock. Recent studies have documented diminished health in IRL dolphins including: high concentrations of mercury [38], lingual and genital papillomas [39], and skin disease (lacaziosis) [40–41]. Moreover, interactions with both commercial and recreational fisheries account for up to 12% of the annual mortality [8, 42].

The Marine Mammal Protection Act requires dolphin stock assessment, and abundance estimates that are necessary to manage stocks and to calculate the level of sustainable anthropogenic mortality (PBR). Data from aerial surveys and observations of movements of IRL dolphins through inlets have suggested temporary emigration/immigration may contribute to fluctuating abundance estimates in portions of the lagoon near inlet access, particularly in response to dramatic declines in water temperature [19, 24] which may influence dolphin movement due to thermoregulatory needs and/or prey movements [43–46]. Photo-identification surveys further support transience occurrence, documenting dolphin movements between the northern portion of the Indian River Lagoon (Mosquito Lagoon) and the St. Johns River (Jacksonville Estuarine system stock-JES), [47] with a 13% exchange in the individuals examined (Nekolny et al. 2017-[20]. Furthermore, genetic research suggest that dolphins inhabiting the Mosquito Lagoon sub-basin may be a disjunct community from the IRL proper as these animals are genetically distinct from the rest of the IRL proper and most closely associated with the JES stock [47], suggesting genetic exchange [21–22]. Year-round mark-recapture surveys are necessary to measure the rate of transient occurrence, the rate of temporary emigration of residents, define the best season to measure resident abundance, to estimate dolphin survival and to determine a precise and current estimate of abundance for this dolphin stock. Likewise, current abundance data are needed to assess the impact of recent mortality events on the population and to calculate PBR. The objectives of this study were to utilize dolphin photo-identification surveys and mark-recapture methodology following a Robust Design survey [48] to estimate abundance, survival and temporary emigration of IRL dolphins. A simulation study was conducted prior to survey initiation to validate the study design (S1 Text, S1 Fig). To assess evidence of transient individuals and measure transient rates, a time since first capture parameter was incorporated into survival sub-models. Lastly, in order to facilitate abundance estimation by sub-basin and primary period (season), closed population capture-recapture models were used.

## Methods

### Ethics Statement

Data collection (vessel-based photo-identification surveys of free-ranging bottlenose dolphins) was conducted under permits issued by NOAA Fisheries under General Authorization Letter of Confirmation No.: 16522, 18182, and 20377-01 in tandem with permits issued by Canaveral National Seashore: CANA-2015-SCI-0010 and the U.S. Fish and Wildlife Service: MI-2016-207R which enabled data collection on the protected/privately owned lands including national wildlife refuges (Cape Canaveral National Seashore, Merritt Island National Wildlife Refuge) and otherwise secured properties (Canaveral Air Force Station, Kennedy Space Center). Data collections consisted of field observations and did not involve animal handling, nor were animals harmed during the course of the study. Due to the benign nature of the study, data collection protocols did not require further assessment by an animal ethics committee and did not raise ethical issues.

### Study area

The Indian River Lagoon is a shallow and diverse estuarine system located along the east coast of central Florida that is open to the Atlantic Ocean at four inlets and consists of three interconnected basins: the Indian River, Banana River and Mosquito Lagoon [49–51] (Fig 1). The 902 km^2^ estuary spans 220 km with a width of 0.93 to 9.30 km [23] extending from Ponce Inlet to Jupiter Inlet [51]. Although most of the estuary is shallow (<1 m at high tide), depths of greater than 5 m occur in the dredged basins and channels of the Intracoastal Waterway (ICW) [49], which encompasses approximately 2.2% of the lagoon. To investigate geographical differences in abundance and density (dolphins/km^2^) and to further investigate movement between basins, the three basins of the IRL were divided into four regions (hereafter termed sub-basins) which present different abiotic and biotic characteristics that could indirectly influence dolphin abundance [52–54]. The Banana River (BR) (202 km^2^) and the Mosquito Lagoon (ML) (140 km^2^) included each sub-basin in its entirety (Fig 1). Because of its large north to south extent, the Indian River basin was divided into two sub-basins: the northern Indian River (NIR) (378 km^2^), previously defined as north of Eau Gallie Causeway [55], with little tidal and non-tidal flushing [52] and the southern Indian River (SIR) (182 km^2^) which consisted of three previously defined basins [55] and includes three of the four inlets (Fig 1). Due to the lack of tidal flushing, the BR and NIR have decreased water quality compared to the majority of ML and SIR [52–54]. Portioning the lagoon into geographically separated basins also allowed comparisons with prior abundance studies [19, 24] and evaluation of previously described communities that inhabit the basins [56].

**Fig 1.**
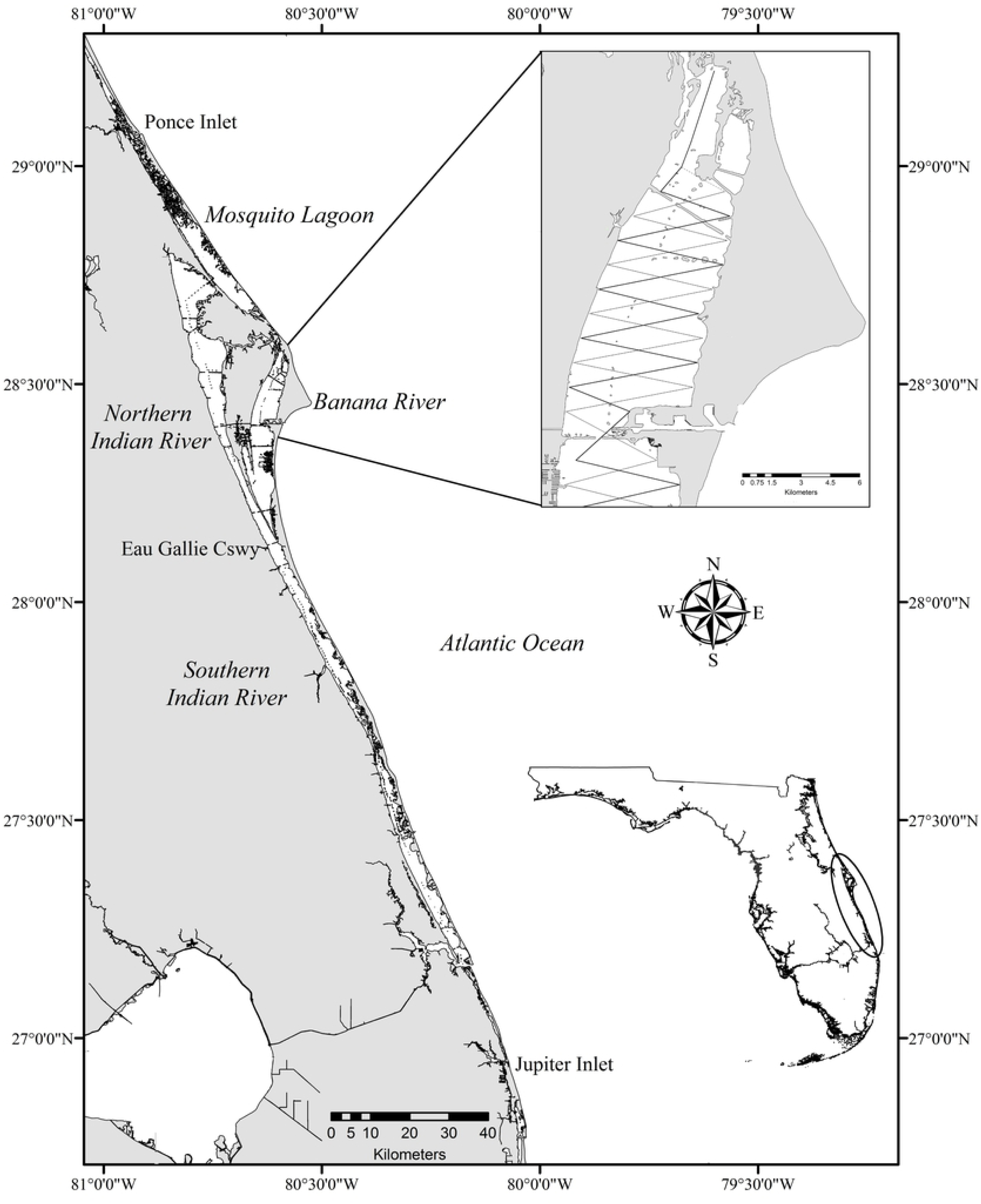
Map of the study area (Indian River Lagoon) along the east coast of Florida (ellipse). Contour lines and alternating saw-tooth transects were utilized throughout the lagoon and are illustrated in the inset of the northern portion of the Banana River. The study area (Ponce Inlet to Jupiter Inlet) was divided into four sub-basins to further evaluate abundance and distribution (Mosquito Lagoon, Banana River, northern Indian River and the southern Indian River).

### Capture-recapture photo-identification surveys

The Robust Design [25, 57] is the preferred method for estimating estuarine dolphin abundance [12–13, 48]. This capture-recapture model is designed to make repeated samples over short temporal periods (secondary sessions) during which population closure is assumed (no emigration/immigration, births or deaths). Sets of replicate secondary samples are repeated over longer periods of time (primary periods) between which the population is considered open. Vessel based capture-recapture surveys were conducted in the Indian River Lagoon between August 2016 and May 2017 and followed a robust survey design which made the following assumptions: population closure (within a primary period) and emigration between sampling being temporary, unique marks that were permanent and correctly read, capture probability was equal during a sampling event, and survival probability was equal among individuals within primary periods [25, 57–58]. The assumption that temporary emigration was temporary was relaxed in some models by including the transient parameter. Additional assumptions required for estimating the proportion of marked individuals (used to adjust for unmarked dolphin abundance) were that marked and unmarked animals did not differ in detection, movement, or survival parameters, and that marked and unmarked individuals mixed randomly [25, 57–58]. Four primary periods (seasons: summer = June-August, fall = September-November, winter = December-February, and spring = March-May) [59] were completed. Primary periods contained three secondary sessions that were completed under optimal conditions (Beaufort Sea State ≤3), in the shortest time period to meet the assumption of closure (target: ≤3 weeks). Secondary sessions (complete survey of the IRL) were separated by at least one day to allow for population mixing [13]. Existing dorsal fin catalogues were utilized and established protocols were closely followed [48]. The survey design used both depth contour lines and alternating saw-tooth transects (total length ∼ 743 km) to minimize capture heterogeneity by providing a broad geographic coverage which allowed individuals to have an equal opportunity of capture (Fig 1). Alternating saw tooth transects (2.5 km apart) were traversed in areas in which the east to west width of the lagoon exceeded 1.25 km. This width was utilized based on a prior study that determined an observation strip, 1.25 km on either side of the survey route, where 95% of sightings occurred [60]. Transects were designed to provide ample coverage of the width of the lagoon (which extends up to 9.3 km) and enabled near complete coverage of the estuary system. Predetermined routes were downloaded into a Global Positioning System unit to ensure transect adherence. Using 11-13 vessels, the entire range for the Indian River Lagoon Estuarine System bottlenose dolphin stock (Ponce Inlet-Jupiter Inlet) [61] was surveyed within one to three days (Fig 1).

Center-console outboard powered vessels ranging from 5-7 m in length, staffed with three to five researchers (driver and left/right observers), were motored (10-12 knots; lowered for slow speed zones and visibility) along a predetermined track in search of dolphins. A sighting was recorded when a dolphin or group of dolphins was observed during survey effort. The vessel was slowed, idled and stopped to photograph dorsal fin markings. Utilizing a Canon EOS digital camera with a 100-400 mm telephoto lens (Canon USA, Inc., Melville, NY, USA), an attempt was made to photograph all dolphins in the group regardless of distinctiveness. A dolphin group was defined as all dolphins within <100 m that were engaged in similar behavior with the same general heading [62]. Calves were defined as swimming in echelon position (very close proximity to the adult’s mid-lateral flank) and <75% of the proximate adult’s size, while young of the year (YOY, less than one year old) were identified based on a body size less than half the adult and swimming in echelon position, also exhibiting some or all of the following: dark color, floppy dorsal fin, presence of fetal lines, extreme buoyancy, and rostrum-first surfacing [63–65]. A datasheet was completed for each sighting and included: time and location parameters-GPS waypoints; latitude and longitude in decimal degrees, behavior, water depth, group size (minimum, maximum, and best estimate) and composition, environmental covariates, and an overall assessment of sighting conditions based on Beaufort sea state, chop height and glare.

### Photo-identification analyses

Identifying and matching individual dorsal fins is critical as capture-recapture methods require the correct identification. Dorsal fin image analyses followed established protocols [66]. Briefly, marked dorsal fins were sorted by notch patterns, with the best photograph serving as the ‘type photograph’ for each dolphin. Subsequently, unambiguous matches with this photograph were accepted as re-identifications if a minimum of two experienced personnel were in agreement of the match. If a distinctly marked dolphin could not be matched to an existing type photograph, it was added to the catalog as a new individual. In order to minimize false matches, images were graded for photographic quality based on a weighed scale of five characteristics [67]. Dorsal fin photographs were then assigned a quality score as follows: Q1=excellent; Q2=average; Q3=poor. The distinctiveness of each dorsal fin was assigned a rating as follows: D1 - very distinctive, D2 - moderately distinct, at least two features or one major feature and D3 - not distinct, few to no features [67–68]. Only Q1/Q2 quality photographs of distinct (D1/D2) dolphins were used in estimating parameters in capture-recapture analyses and similar quality D3 animals provided supplemental information used to adjust abundance estimates for the proportion of unmarked dolphins. Marked (D1-D2) calves and unmarked calves and young of the year were excluded from capture-recapture analyses as they could introduce some non-independence in capture probabilities due to close associations with their mothers [48] and violated the model assumption of independent survival.

### Movements between sub-basins and discovery curve

To evaluate movement between sub-basins, sighting histories for marked dolphins were compiled and movement between the four sub-basins was evaluated. Likewise to further evaluate population closure and site fidelity/temporary emigration the number of times each distinct dolphin was sighted was evaluated and a “discovery curve” [69] was plotted based on the cumulative total number of marked (D1/D2) dolphins across each secondary session.

### Robust Design modeling and model selection

Robust Design capture-recapture models were fit using program MARK [70] via package RMark [71] in R [72]. Parameters estimated included marked dolphin abundance in each primary period (*N*), the probability of apparent survival (φ), the probability of detection (p), and the probability of temporary emigration defined as the probability of an animal being temporarily unavailable for capture if the individual was available during the previous primary period (ϒ’’) or unavailable (ϒ’). Some models also included a time since initial capture (“transient”) parameter to estimate transient presence (animals that are only available for a single detection) [73]. A total of 36 models were fit with combinations of all structural covariates for detection, survival, and temporary emigration parameters. Since capture probability may be influenced by environmental conditions that change over time, detection models allowed detection to vary between primary periods (season), secondary sessions (session), or to be equal across all sampling occasions (.). Because there was no reason to believe that dolphins would respond to our unobtrusive observations (become trap shy), the probability of first detection (p) and the probability of recapture (c) were assumed equal for all models (no behavioral response). Dolphin survival can vary by age class [74]; therefore, to ensure equal survival probabilities, only adults were utilized in analyses. Survival was modeled to vary by primary period (season) or as a constant. Robust Design capture-recapture models are not able to discern true mortality from permanent emigration and thus failure to account for transient animals (those that leave the area after one primary period with no probability of being recaptured) would result in negatively biased estimates of survival [75]. To assess evidence and account for transient individuals a time since initial capture “transient” parameter was included in some models to allow survival during the first primary period after an individual’s initial capture (φ 1) to be estimated separately from its survival later [76]. The theory behind including this parameter is that transients are likely to be sighted in only one primary period and the set of animals captured for the first time in any primary period are a mixture of transients and residents. Therefore, survival estimates for the first primary period after initial capture (φ 1) will include apparent mortality (due to permanent emigration) of the transients and are thereby biased low. The survival estimates for the remaining observations (φ 2) do not include transients and more accurately reflect resident survival. The proportion of transients among the newly-marked animals was calculated as τi= φ i2/ φ i1 and the proportion of transients for the population as Ti = Ni/(Ni + mi) where Ni = the number of newly-marked individuals and mi = the number of previously marked individuals captured within time period I [75, 77]. Models considered allowed transient survival to be constant (φ(transient)) or to vary by season (φ(transient*season). Temporary emigration was modeled to allow Markovian movement in which the probability of availability was dependent on the previous state (available or unavailable) (ϒ’’(.) ϒ’(.)), random movement in which availability probability did not differ based on the previous state (ϒ’’= ϒ’(.)), or no movement models (ϒ’’=1, ϒ’=0). For Markovian and random movement models, the temporary emigration parameters were constrained to be constant throughout the study (ϒ’’(.) =ϒ’(.); ϒ’’= ϒ’(.)). Attempts were made to model temporary emigration as varying over time, but unfortunately parameters were not identifiable, likely due to the reduced number of primary periods (*n* = 4).

Model fit was evaluated using Fletcher’s generic goodness-of-fit statistic (c-hat) [78] calculated by program MARK for the two most parameterized models that included both time (season) and “transient” (time since initial capture) effects on survival, time-specific (season and session) detection parameters, and random temporary emigration parameters. Data were also evaluated pooled across secondary sessions using program RELEASE [79] implemented in Rmark; this test is commonly considered for Robust Design data since the structure of the data collapsed within primary periods is similar to a Jolly-Seber model.

Model comparisons were based on the small-sample adjusted Akaike Information Criterion AICc; [80], calculated with the theoretical number of estimable model parameters (i.e., the column rank of the design matrix), rather than the estimated rank of the variance–covariance matrix, to guard against overfitting the data. Models with ΔAICc < 10 were evaluated for evidence of numerical estimation errors (indicating lack of parameter identifiability). Akaike weights were reported and convey the relative support (compared to all candidate models) for each model on a scale of zero to one [80]. Akaike weights were combined across similar movement model types to evaluate the relative cumulative importance of movement parameters within models [80]. Final abundance estimation was based on model averaging with models that did not show signs of numerical estimation problems. Model average estimates were calculated separately for each of three temporary emigration models.

The proportion of marked individuals within groups was calculated as the number of individuals within groups that were marked (D1 or D2) divided by the total number of adult individuals (D1, D2, and D3). Final abundance estimates were obtained by adjusting the estimated abundance of marked individuals, obtained from capture -recapture models, for the proportion of marked to unmarked animals observed during each primary period. This was obtained by calculating the marked ratio for each group encountered. The final abundance estimates were then calculated as: Ntotal = Nmarked / Proportion Marked. Adjusted standard errors and 95% confidence intervals were calculated following standard methods.

### Abundance by sub-basin

The sub-basins of the IRL present differences which may influence dolphin abundance [52–54], and evidence has suggested differentiation in the dolphin communities by sub-basin including Mosquito Lagoon being a disjunct community from the IRL proper [21–22, 56]. Prior unusual mortality events have been concentrated in small geographic regions within the northern IRL (primarily NIR, BR) [35, 81], therefore abundance estimates by sub-basin imperative to better understand community impacts. Temporal variation in ecological conditions within the lagoon may also influence dolphin abundance [19, 24] making it further desirable to have estimates partitioned into primary periods. Unfortunately, multi-state robust design capture-recapture models which could provide sub-basin-specific abundance estimates require many additional parameters which can introduce numerical difficulties when data are limited to four primary periods. As anticipated, attempts to utilize multi-state robust design capture-recapture models to estimate abundance by sub-basin and primary period resulted in numerical difficulties requiring different methodology. As an alternative, abundance was estimated using closed population capture-recapture abundance models by each of the 16 sub-basin and primary period (season) combinations. Closed population capture-recapture abundance models made the following assumptions: population closure (no births, deaths, immigration or emigration), unique marks that were permanent and correctly read, and equal capture probability for marked and unmarked animals (random mixing after first capture). [82–83]. Closed population capture-recapture models were fit using program MARK [70] via package RMark [71] in R [72]. For each sub-basin and primary period (season), a single model was fit which allowed each secondary session to have a separate capture probability [84]. This model produced reasonable parameter estimates with good precision and provided baseline estimates for comparison.

## Results

### Field effort and photo-identification

From August 2016 through May 2017, 135 boat-based photo-identification surveys (25 survey days) were conducted throughout the Indian River Lagoon to complete four capture-recapture primary periods (12 secondary sessions) for the IRL Estuarine System stock. Each secondary session (complete IRL survey replicate) was completed using 11-13 vessels (11.42 ± 1.00 SD) over a time period of 1-3 d (2.3 ± 0.65 SD) and consisted of 1-3 survey days (2.0 ± 0.43 SD). Each primary period was completed in 13-36 d (20.0 ± 10.9 SD). Surveys ranged from 3.28 -14.25 h (8.72 ± 2.21 SD; total field hours: 1,177.58 h). Over 159,000 photographs were taken from 1,465 dolphin sightings. A total of 4.4% of images of individual dorsal fins were determined to be poor quality (Q3) and were excluded from further analyses. The remaining images were quality one (82.3%) or quality two (8.4%). Sightings were comprised of 5,246 dolphins which consisted of 4,239 adults, 863 calves, and 144 YOYs (Table 1). Dolphin sightings per survey ranged from 0-40 (11.8 ± 7.0 SD) per vessel with 122.08 (± 26.71 SD) sightings per complete IRL survey. A mean of 38.9 dolphins (± 27.26 SD) were sighted on each survey (range: 0-149 animals) with 437.2 dolphins sighted per complete IRL survey (± 101.8 SD). Mean group size was 4.08 (± 4.24 SD; range 1-39) and the largest mean group size was in the summer season (5.08 ± 5.10 SD) (Table 2). Calves and YOYs comprised 19.2% of all animals sighted (Table 1).

**Table 1.** Summary of sightings (dolphin groups) and individual dolphins photographed (unmarked and marked).

| Sub-basin | Dolphin Sightings | Summer<br>Aug 10- 23, 2016 | Fall<br>Nov 1- 15, 2016 | Winter<br>Jan 18 -Feb 3, 2017 | Spring<br>Apr 17-May 23, 2017 | Total |
| --- | --- | --- | --- | --- | --- | --- |
| Banana River | Sightings | 81 | 90 | 99 | 89 | 359 |
|  | Adults/calves/YOYs | 381/77/18 | 213/54/6 | 365/102/8 | 266/72/2 | 1225/305/34 |
|  | Total dolphins | 476 | 273 | 475 | 340 | 1564 |
| Mosquito Lagoon | Sightings | 128 | 87 | 111 | 83 | 409 |
|  | Adults/calves/YOYs | 318/75/9 | 194/39/10 | 212/54/9 | 180/40/6 | 904/208/34 |
|  | Total dolphins | 402 | 243 | 275 | 226 | 1146 |
| Northern Indian River | Sightings | 79 | 60 | 119 | 85 | 343 |
|  | Adults/calves/YOYs | 335/39/10 | 179/26/6 | 305/48/4 | 313/29/8 | 1132/142/28 |
|  | Total dolphins | 384 | 211 | 357 | 350 | 1302 |
| Southern Indian River | Sightings | 65 | 86 | 132 | 71 | 354 |
|  | Adults/calves/YOYs | 200/30/14 | 243/64/14 | 344/81/11 | 191/33/9 | 978/208/48 |
|  | Total dolphins | 244 | 321 | 436 | 233 | 1234 |
| All sub-basins | Sightings | 353 | 323 | 461 | 328 | 1465 |
|  | Adults/calves/YOYs | 1234/221/51 | 829/183/36 | 1226/285/32 | 950/174/25 | 4239/863/144 |
|  | Total dolphins | 1506 | 1048 | 1543 | 1149 | 5246 |

Results are given by age class (adults, calves, young of year-YOYs), primary period and sub-basin. \*Extensive efforts were made to identify both distinctive and non-distinctive dolphins; however, non-distinctive animals may be represented more than once.

**Table 2.** Mean group size (± SD) of Indian River Lagoon dolphins per primary period and sub-basin.

| Primary period | Mosquito Lagoon | Banana River | Northern Indian River | Southern Indian River | Indian River Lagoon |
| --- | --- | --- | --- | --- | --- |
| Summer | 4.27 ± 4.85 | 6.58 ± 6.05 | 5.39 ± 5.13 | 4.40 ± 3.70 | 5.08 ± 5.10 |
| Fall | 3.49 ± 3.43 | 3.46 ± 3.04 | 3.83 ± 4.28 | 4.01 ± 5.04 | 3.68 ± 3.98 |
| Winter | 2.95 ± 2.89 | 5.26 ± 5.61 | 3.21 ± 3.21 | 3.55 ± 2.58 | 3.69 ± 3.73 |
| Spring | 3.01 ± 3.28 | 4.52 ± 5.00 | 4.33 ± 4.19 | 3.83 ± 2.74 | 3.94 ± 3.98 |
| Mean | 3.49 ± 3.81 | 4.92 ± 5.15 | 4.10 ± 4.21 | 3.88 ± 3.56 | 4.08 ± 4.24 |

### Discovery curve and distribution patterns

During the study, 503 distinctively marked individuals were recorded. A greater number of distinctively marked animals were seen in the southern Indian River than in the other sub-basins. A total of 369 marked animals (73.4%) were observed in just one sub-basin, with 124 (33.6%) in the southern Indian River, 115 (31.2%) in Mosquito Lagoon, 73 (19.8%) in the northern Indian River and 57 (15.4%) in the Banana River. A total of 112 marked individuals (22%) were seen in two sub-basins, and 22 (4%) in three sub-basins (NIR, BR and SIR). Of the individuals seen in two sub-basins, the greatest exchange was seen between the northern Indian River and the Banana River, followed by the northern Indian River and the southern Indian River (Fig 2). Eighty-seven percent of the distinct individuals in Mosquito Lagoon were only observed in Mosquito Lagoon, while the remaining 13% were also observed in the northern Indian River (Fig 2). Movement of distinctly marked animals between the Banana River and all of the other sub-basins was observed, excluding Mosquito Lagoon (Fig 2). Exchange between the northern Indian River and all other basins were observed (Fig 2). Distinct individuals in the southern Indian River moved into all other basins except for Mosquito Lagoon (Fig 2). Of the 503 distinctly marked animals, 84 animals (16.7%) were sighted during a single survey only (Fig 3) and were treated as transients in some Robust Design models. The 84 animals were distributed across all primary periods (fall= 17; spring = 18, summer = 29, and winter = 20) and sub-basins (Mosquito Lagoon = 23, Banana River =17, northern Indian River=22, southern Indian River = 22). Marked animals were sighted in one (*n* = 105; 20.9%), two (*n* = 155, 30.8%), three (*n* = 168, 33.4%) or four seasons (*n* = 75, 14.9%). The sighting frequency for marked individuals ranged from one and nine secondary sessions (Fig 3). The discovery rate for new marked animals increased steeply during the first primary period (summer) and then rose steadily throughout the study until reaching a plateau in the final secondary session (Fig 4).

**Fig 2.**
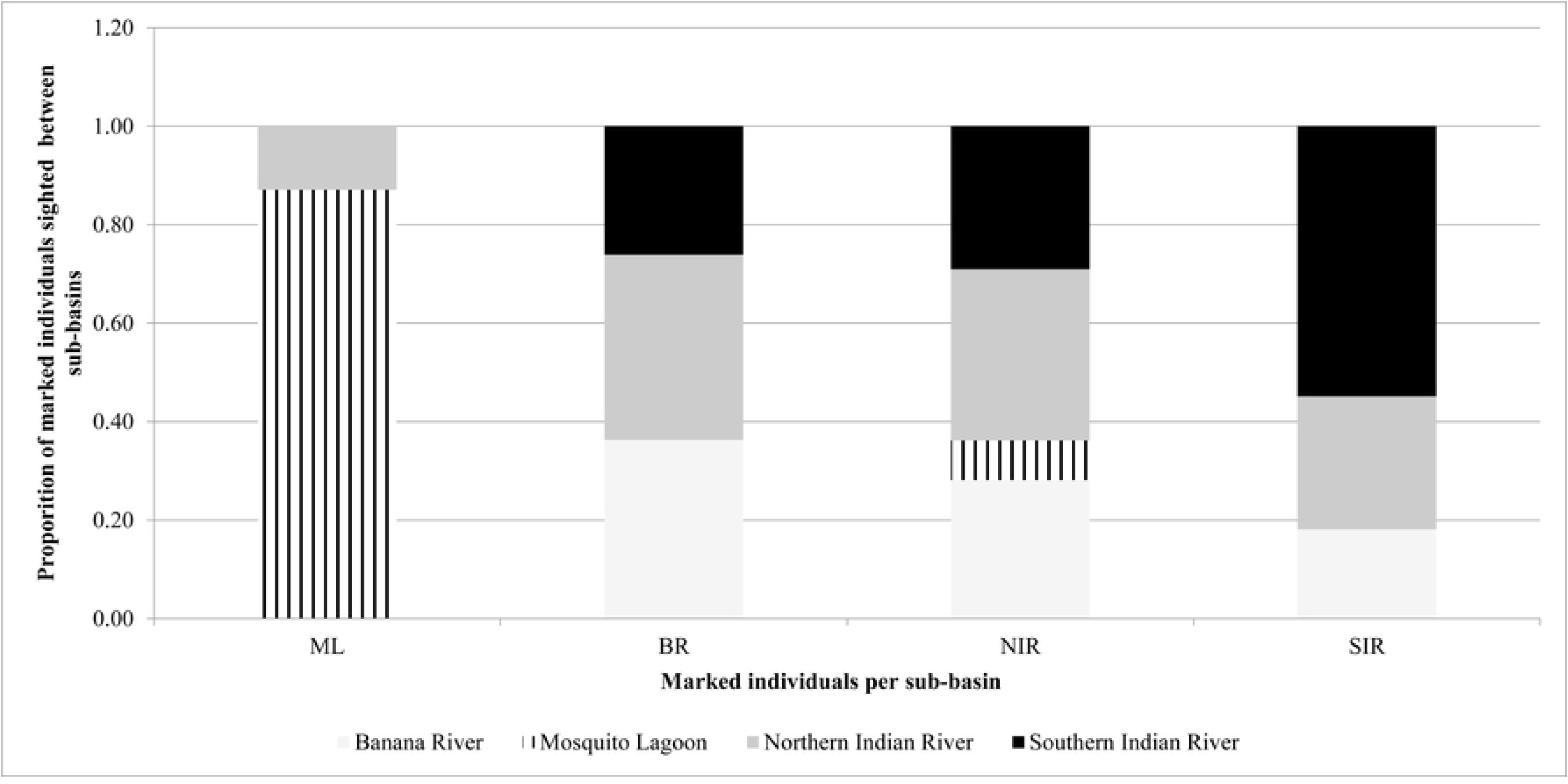
The proportion of exchange of marked individuals sighted between sub-basins. Individuals were either exclusively sighted in one sub-basin or were sighted in additional sub-basins. Sub-basins included: ML= Mosquito Lagoon, BR= Banana River, NIR= northern Indian River, SIR =southern Indian River.

**Fig 3.**
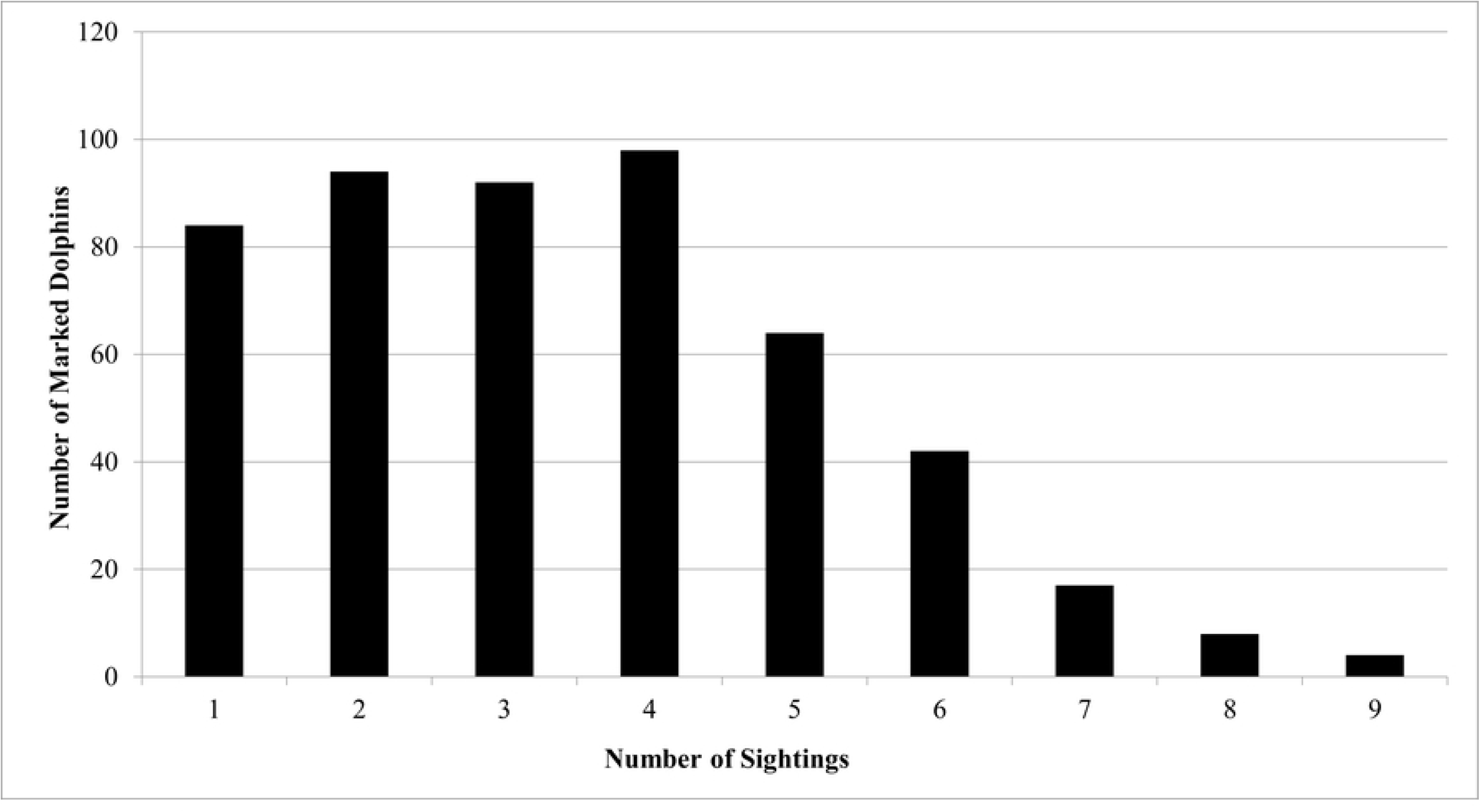
Sighting frequency of marked bottlenose dolphins from photo-identification surveys.

**Fig 4.**
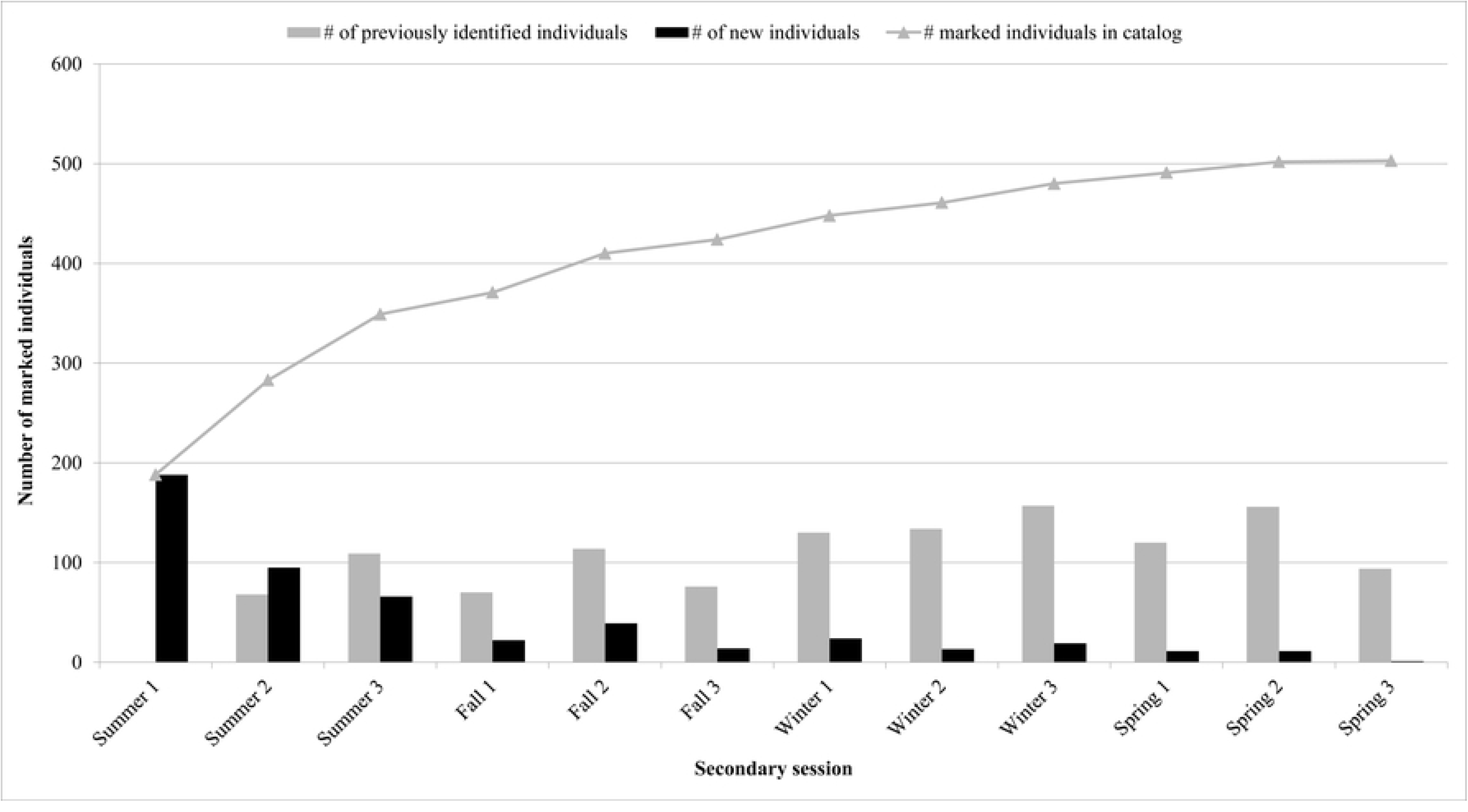
Number of marked individuals sighted during Indian River Lagoon dolphin photo-identification surveys and discovery curve.

### Robust Design modeling and model selection

Fletcher c-hat was 1.03 for both of the two most parameterized models indicating the data did not have a significant degree of overdispersion. The goodness-of-fit test implemented in program RELEASE also indicated no evidence for lack of fit (Chi-square = 78.7, df = 61, p = 0.06). Since there was no evidence for overdispersion for the general data structure or the most parametrized models, models were not adjusted for overdispersion (c-hat). Twelve of the 36 models were clearly supported as being superior (Table 3, S1 Table). Of these, five had ΔAICc < 2 indicating a high level of support, and the remaining seven had 2 < ΔAICc < 10 indicating a moderate level of support [85]. All of the best supported models included parameters to estimate a detection rate for each secondary session. The best supported model allowed survival to vary by primary period (season) and by time since initial capture (“transient”) and included the no movement model of temporary emigration (low level emigration). The next best supported model also allowed survival to vary by time since initial capture but had random temporary emigration constrained to be equal across seasons. Model average estimates of detectability ranged from 0.20 to 0.42 between secondary sessions (Table 4, Fig 5). The model average estimates of seasonal (three month) marked dolphin survival for residents only (sighted in two or more seasons) ranged from 0.85-1.00 (Table 5). Parameter estimates for each well-supported model are reported (S1 Table).

**Fig 5.**
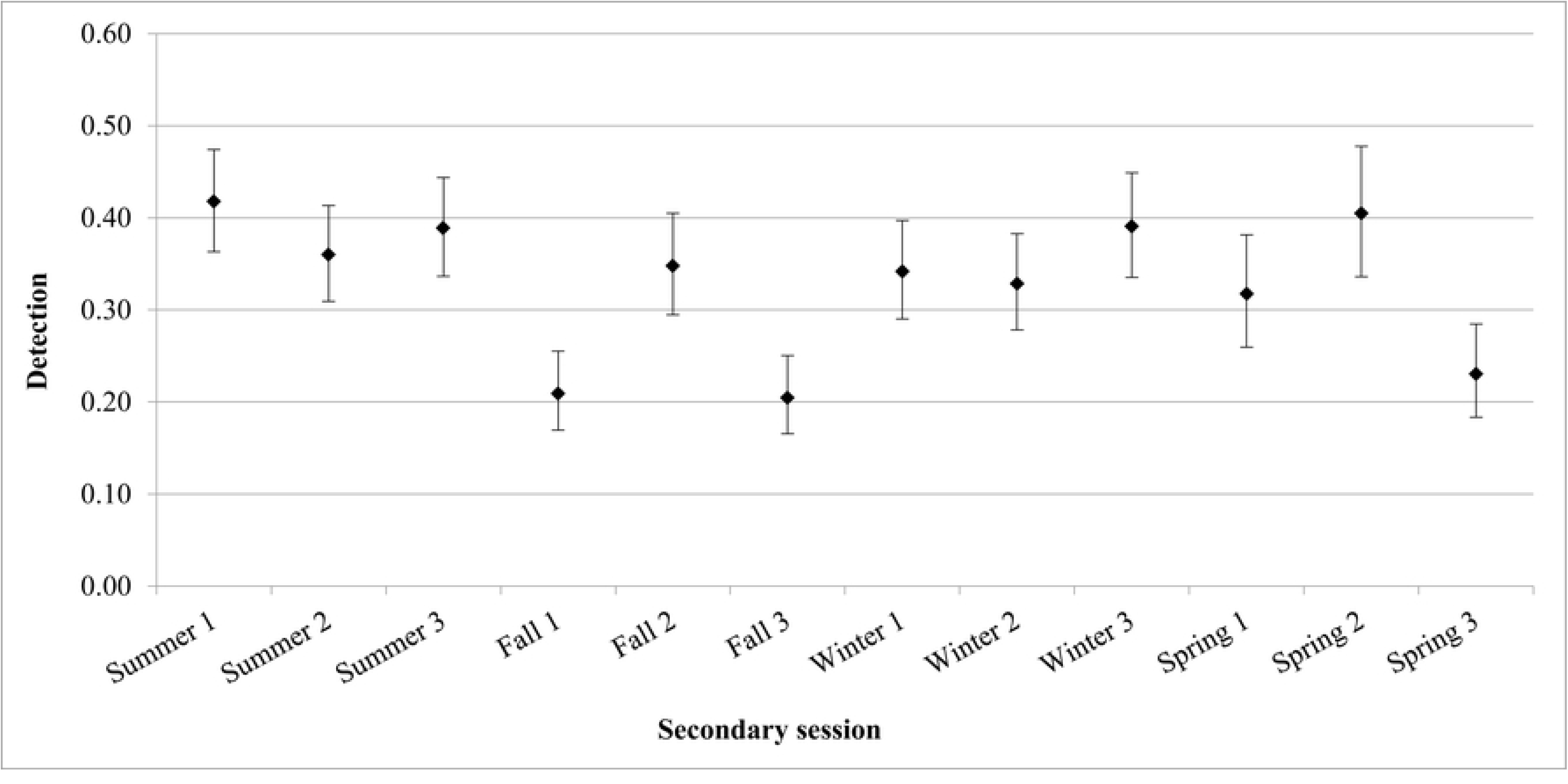
Model averaged estimates of bottlenose dolphin detection (95%CI) by secondary session. Detection was calculated using model averaged Robust Design models for capture-recapture study of the Indian River Lagoon Estuarine System stock of bottlenose dolphin during 2016-2017.

**Table 3.** Results of model selection for Robust Design models.

| Model | Temporay<br>emigration<br>model | AICc | $\Delta AICc$ | #<br>parameters | Deviance | Weight |
| --- | --- | --- | --- | --- | --- | --- |
| $\phi(\text{transient}*\text{season})Y''=1 \ Y'=0 \ p=c(\text{season}*session)$ | No movement | -5371.38 | 0.00 | 21 | 497.86 | 0.19 |
| $\phi(\text{transient})Y''=Y'(\cdot) \ p=c(\text{season}*session)$ | Random | -5370.59 | 0.79 | 19 | 502.75 | 0.13 |
| $\phi(\cdot)Y''=Y'(\cdot) \ p=c(\text{season}*session)$ | Random | -5370.39 | 0.99 | 18 | 504.98 | 0.12 |
| $\phi(\text{transient}*season)Y''=Y'(\cdot) \ p=c(\text{season}*session)$ | Random | -5370.29 | 1.09 | 22 | 496.90 | 0.11 |
| $\phi(\text{season})Y''=1 \ Y'=0 \ p=c(\text{season}*session)$ | No movement | -5369.72 | 1.66 | 19 | 503.62 | 0.08 |
| $\phi(\text{transient})Y''(\cdot)Y'(\cdot) \ p=c(\text{season}*session)$ | Markovian | -5369.35 | 2.03 | 20 | 501.93 | 0.07 |
| $\phi(\cdot)Y''(\cdot)Y'(\cdot) \ p=c(\text{season}*session)$ | Markovian | -5369.17 | 2.21 | 19 | 504.16 | 0.06 |
| $\phi(\text{season})Y''=Y'(\cdot) \ p=c(\text{season}*session)$ | Random | -5369.07 | 2.31 | 20 | 502.21 | 0.06 |
| $\phi(\text{transient}*season)Y''(\cdot)Y'(\cdot) \ p=c(\text{season}*session)$ | Markovian | -5368.52 | 2.86 | 23 | 496.61 | 0.05 |
| $\phi(\cdot)Y''=1 \ Y'=0 \ p=c(\text{season}*session)$ | No movement | -5368.44 | 2.93 | 17 | 508.98 | 0.04 |
| $\phi(\text{season})Y''(\cdot)Y'(\cdot) \ p=c(\text{season}*session)$ | Markovian | -5368.14 | 3.24 | 21 | 501.10 | 0.04 |
| $\phi(\text{transient})Y''=1 \ Y'=0 \ p=c(\text{season}*session)$ | No movement | -5367.91 | 3.47 | 18 | 507.47 | 0.03 |
| $\phi(\text{transient}*season)Y''=1 \ Y'=0 \ p=c(session)$ | No movement | -5321.20 | 50.18 | 13 | 564.37 | 0.00 |
| $\phi(\text{transient})Y''=Y'(\cdot) \ p=c(session)$ | Random | -5320.65 | 50.73 | 11 | 568.98 | 0.00 |
| $\phi(\cdot)Y''=Y'(\cdot) \ p=c(session)$ | Random | -5320.41 | 50.97 | 10 | 571.24 | 0.00 |
| $\phi(\text{transient}*season)Y''=Y'(\cdot) \ p=c(session)$ | Random | -5319.73 | 51.65 | 14 | 563.80 | 0.00 |
| $\phi(\cdot)Y''=1 \ Y'=0 \ p=c(session)$ | No movement | -5319.72 | 51.66 | 9 | 573.95 | 0.00 |
| $\phi(\text{season})Y''=1 \ Y'=0 \ p=c(session)$ | No movement | -5319.64 | 51.74 | 11 | 569.98 | 0.00 |
| $\phi(\text{transient})Y''(\cdot)Y'(\cdot) \ p=c(session)$ | Markovian | -5319.45 | 51.93 | 12 | 568.15 | 0.00 |
| $\phi(\text{transient})Y''=1 \ Y'=0 \ p=c(session)$ | No movement | -5319.35 | 52.03 | 10 | 572.30 | 0.00 |
| $\phi(\cdot)Y''(\cdot)Y'(\cdot) \ p=c(session)$ | Markovian | -5319.28 | 52.10 | 11 | 570.34 | 0.00 |
| $\phi(\text{season})Y''=Y'(\cdot) \ p=c(session)$ | Random | -5318.50 | 52.88 | 12 | 569.09 | 0.00 |
| $\phi(\text{transient}*season)Y''(\cdot)Y'(\cdot) \ p=c(session)$ | Markovian | -5317.99 | 53.39 | 15 | 563.51 | 0.00 |
| $\phi(\text{season})Y''(\cdot)Y'(\cdot) \ p=c(session)$ | Markovian | -5317.53 | 53.85 | 13 | 568.03 | 0.00 |
| $\phi(\text{season})Y''=Y'(\cdot) \ p=c(\cdot)$ | Random | -5291.59 | 79.79 | 9 | 602.08 | 0.00 |
| $\phi(\text{transient})Y''=Y'(\cdot) \ p=c(\cdot)$ | Random | -5291.47 | 79.91 | 8 | 604.23 | 0.00 |
| $\phi(\text{transient}*season)Y''=Y'(\cdot) \ p=c(\cdot)$ | Random | -5290.74 | 80.64 | 11 | 598.88 | 0.00 |
| $\phi(\text{season})Y''(\cdot)Y'(\cdot) \ p=c(\cdot)$ | Markovian | -5289.57 | 81.81 | 10 | 602.08 | 0.00 |
| $\phi(\text{transient})Y''(\cdot)Y'(\cdot) \ p=c(\cdot)$ | Markovian | -5289.47 | 81.91 | 9 | 604.21 | 0.00 |
| $\phi(\cdot)Y''=Y'(\cdot) \ p=c(\cdot)$ | Random | -5289.19 | 82.19 | 7 | 608.52 | 0.00 |
| $\phi(\text{transient}*season)Y''(\cdot)Y'(\cdot) \ p=c(\cdot)$ | Markovian | -5288.96 | 82.42 | 12 | 598.64 | 0.00 |
| $\phi(\cdot)Y''(\cdot)Y'(\cdot) \ p=c(\cdot)$ | Markovian | -5287.21 | 84.17 | 8 | 608.48 | 0.00 |
| $\phi(\text{transient}*season)Y''=1 \ Y'=0 \ p=c(\cdot)$ | No movement | -5285.69 | 85.69 | 10 | 605.96 | 0.00 |
| $\phi(\text{season})Y''=1 \ Y'=0 \ p=c(\cdot)$ | No movement | -5284.13 | 87.25 | 8 | 611.56 | 0.00 |
| $\phi(\text{transient})Y''=1 \ Y'=0 \ p=c(\cdot)$ | No movement | -5278.64 | 92.74 | 7 | 619.08 | 0.00 |
| $\phi(\cdot)Y''=1 \ Y'=0 \ p=c(\cdot)$ | No movement | -5277.79 | 93.59 | 6 | 621.94 | 0.00 |

Parameters for a capture-recapture study of the Indian River Lagoon Estuarine System stock of bottlenose dolphins during four primary periods (2016-2017) include: apparent survival ( $\phi$ ), probability of first detection ( $p$ ), temporary emigration ( $Y'$ ), and the probability of recapture( $c$ ) and were constrained constant ( · ) or allowed to vary by primary period (season) and/or secondary session. A transient parameter (time since first capture) was modeled to assess evidence of
transient individuals. Models allowed apparent transient survival to be constant ( $\phi$ (transient)), or to vary by primary period ( $\phi$ (transient\*season)).

**Table 4.** Model averaged estimates detection (p) by secondary session.

| Session | p | SE | LCL | UCL |
| --- | --- | --- | --- | --- |
| Summer session 1 | 0.42 | 0.03 | 0.36 | 0.47 |
| Summer session 2 | 0.36 | 0.03 | 0.31 | 0.41 |
| Summer session 3 | 0.39 | 0.03 | 0.34 | 0.44 |
| Fall session 1 | 0.21 | 0.02 | 0.17 | 0.26 |
| Fall session 2 | 0.35 | 0.03 | 0.29 | 0.41 |
| Fall session 3 | 0.20 | 0.02 | 0.17 | 0.25 |
| Winter session 1 | 0.34 | 0.03 | 0.29 | 0.40 |
| Winter session 2 | 0.33 | 0.03 | 0.28 | 0.38 |
| Winter session 3 | 0.39 | 0.03 | 0.34 | 0.45 |
| Spring session 1 | 0.32 | 0.03 | 0.26 | 0.38 |
| Spring session 2 | 0.40 | 0.04 | 0.34 | 0.48 |
| Spring session 3 | 0.23 | 0.03 | 0.18 | 0.28 |

Detection was estimated with Robust Design for capture-recapture models for Indian River Lagoon dolphins (2016-2017). \*SE=Standard deviation; LCL=Lower confidence limit; UCL=Upper confidence limit

**Table 5.** Model averaged estimates of adult survival rates (S) (95% CI) between primary periods calculated.

| Survival between primary periods | Random S | SE | No movement S | SE | Markovian S | SE |
| --- | --- | --- | --- | --- | --- | --- |
| S transient Summer-Fall | 0.94 (0.88, 0.97) | 0.02 | 0.93 (0.88, 0.96) | 0.02 | 0.95 (0.83, 0.99) | 0.03 |
| S resident Fall-Winter | 0.98 (0.74, 1.00) | 0.03 | 0.98 (0.74, 1.00) | 0.03 | 1.00 (1.00, 1.00) | 0.00 |
| S transient Fall-Winter | 0.93 (0.74, 0.98) | 0.05 | 0.90 (0.69, 0.98) | 0.06 | 0.94 (0.67, 0.99) | 0.06 |
| S resident Winter-Spring | 0.92 (0.60, 0.99) | 0.08 | 0.85 (0.64, 0.94) | 0.07 | 0.93 (0.53, 0.99) | 0.08 |
| S transient Winter-Spring | 0.90 (0.61, 0.98) | 0.08 | 0.84 (0.57, 0.95) | 0.09 | 0.90 (0.59, 0.98) | 0.08 |

Estimates are calculated by averaging within each of the temporary emigration model types
using Robust Design models for capture-recapture study of the Indian River Lagoon Estuarine
System stock of bottlenose dolphins during 2016-2017. Transient refers to survival estimates for
the first captures (including transients and residents) while resident refers to survival estimates
for all subsequent captures (residents only). \*SE = standard error

No movement and random movement models were a better fit than Markovian models of temporary emigration (Table 3). The combined Akaike model weight for random movement models was 0.42, while no movement models had a cumulative weight of 0.36 and Markovian movement models had a cumulative weight of 0.22. The probability that the dolphin was available (inside the study area) in the prior sampling period and subsequently moved to the unavailable state (outside the study area) in the next sampling period (ϒ’’) was estimated to be quite low with the model average estimate for all models being 0.05 (SE = 0.05). The probability that a dolphin was unavailable for observation (outside the study area) in the prior sampling period and remained unavailable in the next sampling period (ϒ’) was greater, but estimates lacked precision 0.48 (SE = 0.44). The proportion of transients (transient rate) among the marked population was estimated from 0.06 in winter to 0.07 in fall, the only seasons for which the proportion of transients could be estimated [75, 77]. The proportion of marked individuals ranged from 0.40 – 0.45 between primary periods (Table 6). Abundance for the IRL Estuarine System stock ranged from 981 (95% CI: 882-1,090) dolphins in the winter season (primary period) to 1,078 (95% CI: 968-1,201) dolphins in the summer season (primary period) (Table 6, Fig 6). Mean estimated dolphin abundance was 1,032 (95% CI: 969 – 1,098) (Table 6). IRL dolphin density ranged from 1.09-1.20 dolphins/km^2^ (1.15 ± 0.05 SD) (Table 6).

**Fig 6.**
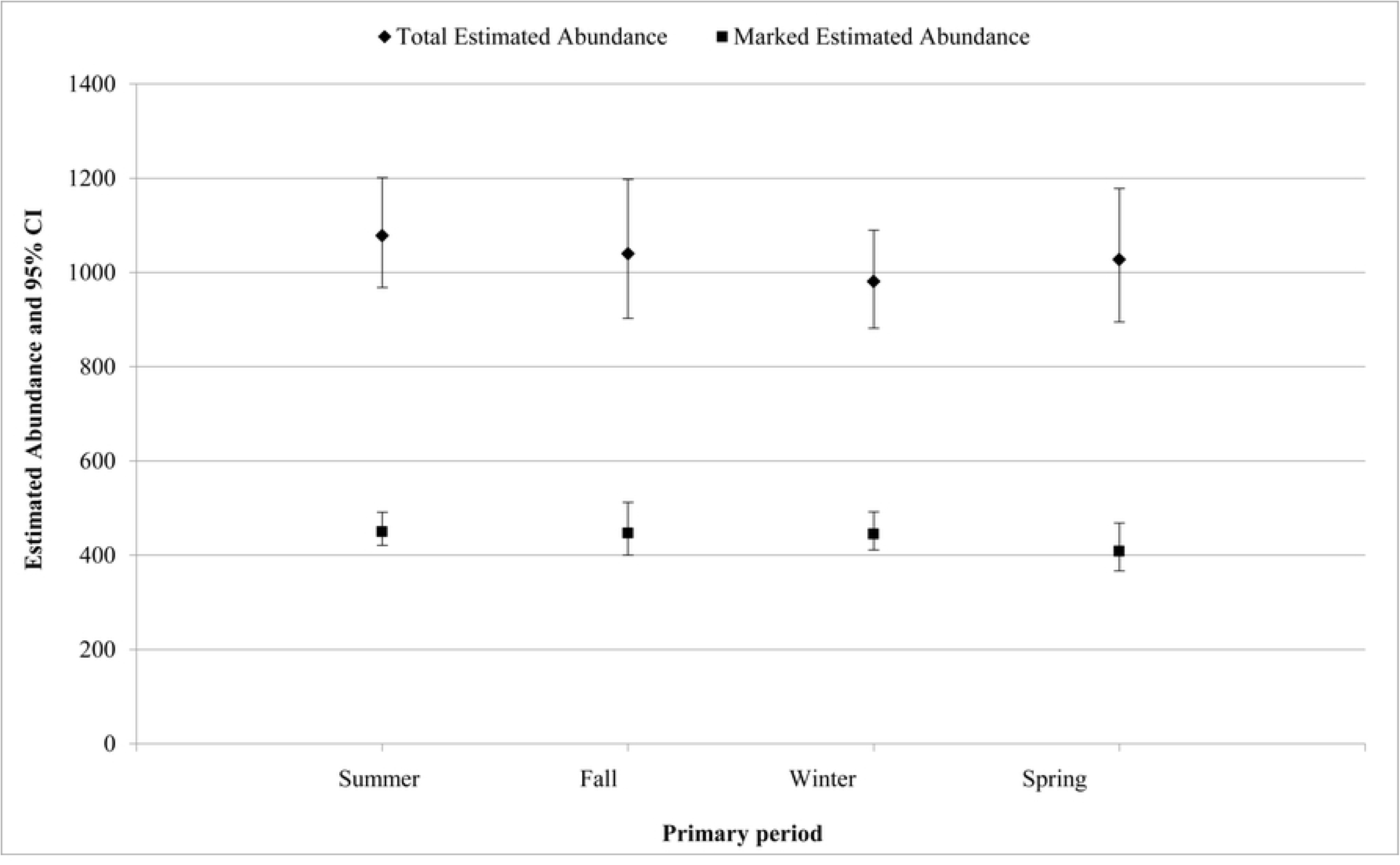
Estimated seasonal abundance (95% CI) for Indian River Lagoon bottlenose dolphins (2016-2017). Abundance was calculated using Robust Design models for capture-recapture study of Indian River Lagoon Estuarine System stock bottlenose dolphins during four primary periods (seasons). Marked abundance estimates include marked animals only; total abundance estimates for the entire population were obtained by adjusting for the ratio of marked: unmarked individuals observed.

**Table 6.** Estimated abundance (95% CI) by primary period for the Indian River Lagoon dolphin population (2016-2017).

| Primary period | *N* (marked) | SE | Proportion marked | Adjusted SE | *N* (marked and unmarked) | Density (dolphins/km<sup>2</sup>) |
| --- | --- | --- | --- | --- | --- | --- |
| Summer | 450 (421, 491) | 18 | 0.42 | 59 | 1078 (968, 1201) | 1.20 |
| Fall | 447 (400, 512) | 28 | 0.43 | 75 | 1040 (903, 1198) | 1.15 |
| Winter | 445 (411, 492) | 20 | 0.45 | 53 | 981 (882, 1090) | 1.09 |
| Spring | 408 (367, 468) | 25 | 0.40 | 72 | 1027 (895, 1178) | 1.14 |

Abundance estimates are given for the marked animals (*N* marked) used in multi-state Robust
Design capture-recapture models, and for the total population (*N* marked and unmarked) which
is obtained by adjusting for the proportion of unmarked animals observed in each primary period
(season). \*SE = standard error

### Abundance by sub-basin

Abundance estimates varied by sub-basin and season (Table 7, Fig 7). The greatest mean abundance was in the southern Indian River (364, 95% CI: 303-438; range: 223-475), followed by the Banana River (349, 95% CI: 296-411; range: 204-482) and northern Indian River (346, 95% CI: 285-420; range: 260-491); while the lowest mean abundance was observed in Mosquito Lagoon (178, 95% CI: 152-209; range: 130-225). The lowest seasonal abundance was observed in Mosquito Lagoon during the fall season (130, 95% CI: 87-194) (Table 7). Dolphin density (dolphins/km^2^) varied by sub-basin with the largest mean density in the southern Indian River (2.00 ± 0.60 SD), followed by the Banana River (1.72 ± 0.57 SD), Mosquito Lagoon (1.23 ± 0.32 SD), and the northern Indian River (0.91 ± 0.27 SD) (Table 7).

**Fig 7.**
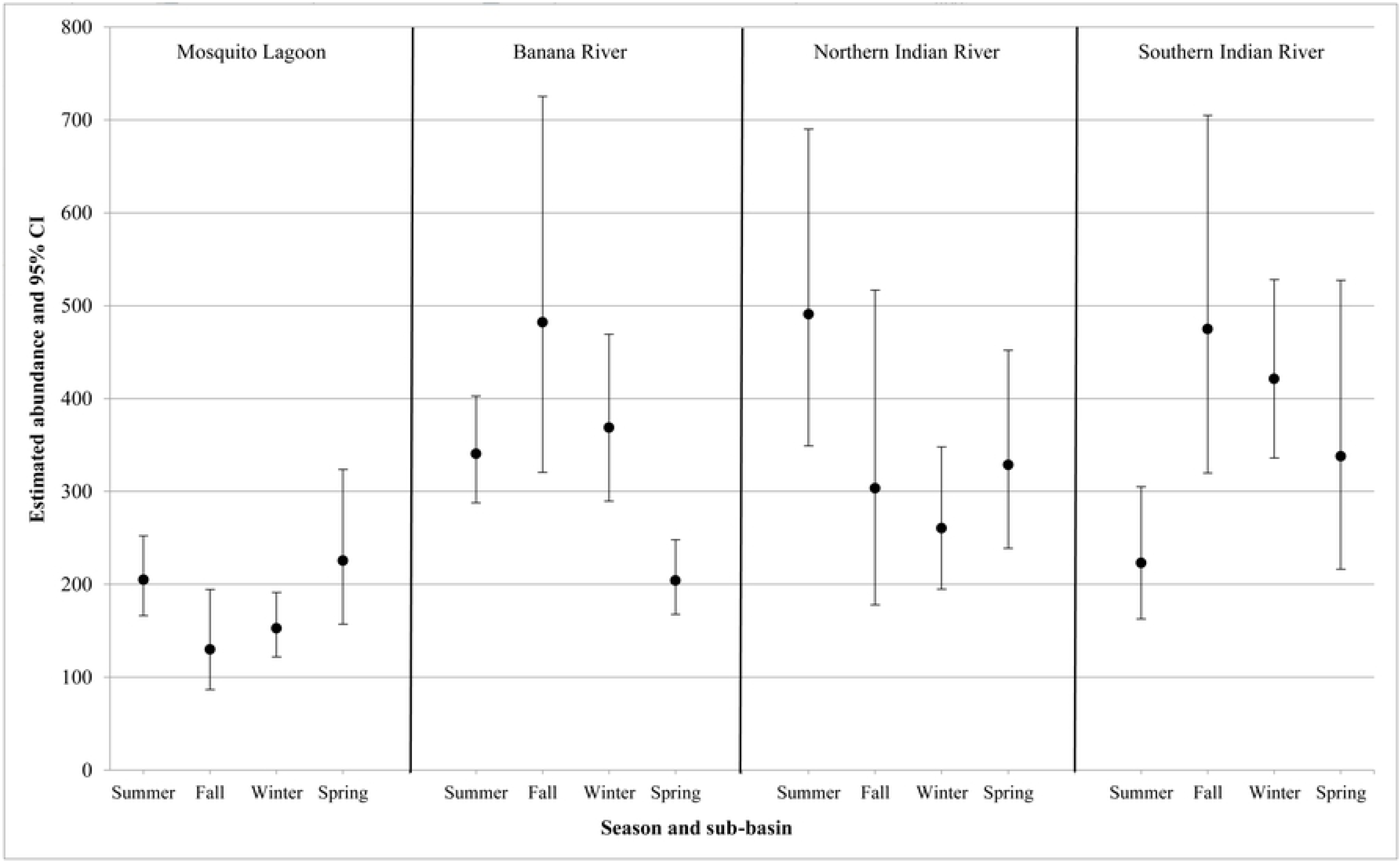
Estimated dolphin abundance calculated using closed population capture-recapture abundance models by sub-basin and primary period. Sub-basins included: ML=Mosquito Lagoon, BR=Banana River, NIR=Northern Indian River, SIR=Southern Indian River. Total dolphin abundance estimates were obtained by adjusting for the ratio of marked: unmarked individuals observed in the Indian River Lagoon (2016-2017).

**Table 7.** Estimated abundance by primary period and sub-basin (95% CI) for Indian River Lagoon dolphins (2016-2017).

| Sub-basin | Primary period | Marked ratio | *N* (marked) | SE | *N* (marked and unmarked) | Adjusted SE | Density (dolphins/km<sup>2</sup>) |
| --- | --- | --- | --- | --- | --- | --- | --- |
| ML | Summer | 0.51 | 105 (91,133) | 10.14 | 205 (166, 252) | 21.85 | 1.46 |
| ML | Fall | 0.41 | 54 (41, 86) | 10.37 | 130 (87, 194) | 26.99 | 0.93 |
| ML | Winter | 0.45 | 69 (61, 89) | 6.55 | 153 (122, 191) | 17.63 | 1.09 |
| ML | Spring | 0.50 | 113 (87, 168) | 19.55 | 225 (157, 324) | 41.98 | 1.61 |
| BR | Summer | 0.33 | 112 (104, 127) | 5.63 | 341 (288, 403) | 29.18 | 1.69 |
| BR | Fall | 0.30 | 143 (107, 216) | 26.49 | 482 (321, 725) | 101.50 | 2.39 |
| BR | Winter | 0.23 | 86 (77, 107) | 7.38 | 369 (290, 469) | 45.58 | 1.82 |
| BR | Spring | 0.32 | 65 (61, 77) | 3.87 | 204 (168, 248) | 20.38 | 1.01 |
| NIR | Summer | 0.39 | 193 (148, 275) | 31.34 | 491 (349, 690) | 85.99 | 1.30 |
| NIR | Fall | 0.36 | 109 (74, 192) | 28.00 | 303 (178, 517) | 83.99 | 0.80 |
| NIR | Winter | 0.36 | 94 (78, 127) | 11.87 | 260 (195, 348) | 38.73 | 0.69 |
| NIR | Spring | 0.41 | 134 (107, 188) | 19.75 | 329 (239, 452) | 53.80 | 0.87 |
| SIR | Summer | 0.66 | 147 (115, 208) | 22.67 | 223 (163, 305) | 35.95 | 1.23 |
| SIR | Fall | 0.57 | 270 (195, 411) | 52.99 | 475 (320, 705) | 96.61 | 2.61 |
| SIR | Winter | 0.57 | 240 (201, 303) | 25.48 | 421 (336, 528) | 48.75 | 2.32 |
| SIR | Spring | 0.56 | 188 (132, 304) | 41.72 | 338 (216, 527) | 77.79 | 1.86 |

Abundance estimates (*N*) calculated using closed capture-recapture models are given for the marked animals, and for the total population which is obtained by adjusting for the proportion of unmarked animals observed in each primary period (season). Sub-basins include the Mosquito Lagoon (ML), Banana River (BR), Northern Indian River (NIR) and southern Indian River (SIR). \*SE = standard error

The mean proportion of marked individuals 43 ± 0.12 SD varied between sub-basins (range: 29.4 -58.8%). The largest mean proportion of marked individuals occurred in the southern Indian River (58.8%), followed by Mosquito Lagoon (47.0%), the Northern Indian River (38.0%) and the lowest proportion of marked individuals occurring in the Banana River (29.4%) (Table 7). The proportion of marked individuals within sub-basins varied slightly between primary periods (seasons) (Table 7).

## Discussion

Recently, much attention has been given to assessing the status of bottlenose dolphin populations in bays, sounds, and estuaries [1–3, 13, 73, 86]. Measuring abundance and demography of these populations is often difficult due to uncertainty about population boundaries [2, 13, 73, 86]. This study is the first to implement a Robust Design capture-recapture design to estimate bottlenose dolphin abundance in the Indian River Lagoon estuary. Due to the predominantly enclosed nature of the IRL, this study was able to implement one of the most comprehensive sampling designs for a dolphin Robust Design study. However achieving this level of coverage required enormous field effort over an expansive and complex geographic area. Estimates obtained from this study achieved greater precision, but were comparable to recent studies of the IRL dolphin abundance estimates [19]; suggesting a stable population. The study provided the first estimates of survival, transient rate, and temporary emigration rates for the IRL dolphin stock, parameters which are essential to management. The strong support for the no movement model and a low estimated transient rate support prior studies of IRL dolphins which have indicated a primarily residential population. While the transient rate was low, separating dolphins sighted only once (transients) and dolphin sighted more than one time more accurately estimated resident IRL dolphin survival rate. Robust Design capture-recapture parameter estimates provide critical guidance to stock managers for a dolphin population with diminished health, where anthropogenic impacts likely exceed the level of sustainable anthropogenic mortality.

The mean dolphin abundance for the Indian River Lagoon Estuarine System stock (1,032; 95% CI: 969-1,098) was similar to estimates from a multi-year line transect aerial surveys (1,032 dolphins; 95% CI: 809 -1,255) [19]. Mean density estimates were also very similar; 1.10 ± 0.46 dolphins/km^2^ from aerial survey data [19] vs. 1.15 ± 0.05 dolphins/km^2^ from the current study. This is interesting since the two studies differed in duration; mark-recapture surveys were conducted over a one year period, while aerial surveys over six years. Stability in the mean annual density and abundance estimates for the IRL Estuarine System stock over these time periods (2006-2011 vs. 2016-2017) is consistent with a stable population. Seasonal fluctuations in the aerial line transect data indicated larger abundance in winter vs. summer [19]. In contrast, mark-recapture surveys indicated only very slight changes in abundance between seasons. Winter aerial surveys were conducted during several unusual cold temperature and hard-freeze events which may have affected the abundance in the Mosquito Lagoon and the southern Indian River [19]. In contrast, photo-identification surveys were conducted over a mild winter and did not measure an increase in winter abundance in Mosquito Lagoon. Seasonal variance in abundance was observed in the southern Indian River (lower summer abundance and increased winter abundance) that was similar to observations from prior studies [19]. Similarly, the greatest mean abundance for a sub-basin was observed in the southern Indian River (364, 95% CI: 303-438), with the lowest abundance in Mosquito Lagoon (178, 95% CI: 152-209) as reported in prior estimates (southern Indian River: 347; 95% CI = 202-492, Mosquito Lagoon: 206; 95% CI = 126 -286) [19]. Estimates obtained from both surveys likely included transient dolphins (dolphins that permanently moved beyond the study area during the study). In the northern end of the study area, dolphins from Mosquito Lagoon have been documented traveling well beyond the boundaries of the IRL into the Atlantic Ocean and adjacent northern estuarine waters, with some animals ranging ∼138 km north into the JES [19–20, 47]. Efforts should be made to resolve stock delineation and community structure to better model movements and the occurrence of transients for future capture-recapture studies.

The successful completion of the project involved an extensive collaborative effort and required the coordination of dozens of experienced personnel and 11-13 vessels traversing ∼728 km of transect-contour lines, to provide near complete coverage of the expansive lagoon. This level of effort is unusual among published efforts to estimate estuarine bottlenose dolphin abundance and represents one of the most comprehensive study designs implemented for bay, sound and estuarine dolphin Robust Design capture-recapture studies. The completion of surveys within brief secondary sessions relied on optimal weather being available throughout each primary period. This was often difficult over the expansive area and involved complicated logistical planning around personnel, equipment, and closure of partial sub-basins due to rocket launch operations in the immediate area. Future studies to estimate dolphin abundance for the Indian River Lagoon Estuarine System stock could reduce survey effort while maintaining precision by utilizing parameter estimates from this study to develop a reduced effort sampling design. The current study produced accurate and precise abundance estimates, however survival estimates were more variable and possibly influenced by the low number of primary periods (*n* = 4). Conducting future survey efforts over a longer time period, with increased numbers of primary periods, could aid in the precision of survival estimates by providing more information for estimating temporal parameters. Exploring the effect of sampling sub-basins rather than near complete coverage of the expansive area, utilization of more complex models (Robust Design spatial capture-recapture), and incorporating detection covariates to further improve precision may help reduce the need for extensive field effort and personnel. Lastly, the northern portion of the study area did not incorporate the full extent of the home range of dolphins inhabiting Mosquito Lagoon, as animals in this region may range well beyond the northern IRL border [20]. This could have potentially caused an availability bias due to temporary emigration. Future studies should consider resolving this issue in stock delineation with comprehensive studies of movement patterns in this area.

A prior study of dolphin communities found two communities occurring in Mosquito Lagoon, however in that study the northernmost community had limited sampling since home ranges for these animals were suspected to extend north of the IRL [56]. Four additional communities were described occurring with one occupying a portion of the NIR and BR, and three occurring in the SIR (as defined in this study), with all communities having a degree of overlap with adjacent basins [56]. Prior research indicated that dolphins sighted in Mosquito Lagoon exhibited a strong site fidelity to that sub-basin (71% exclusive to ML) [87]. Similarly, 87% of dolphins sighted in Mosquito Lagoon during the current short-term study were only observed in this IRL sub-basin. Small, negligible differences between studies may be reflective of the potential for IRL dolphin ranging patterns to expand over time [60] or the short-term nature of the current study. The greatest exchange between sub-basins was observed between the northern Indian River and the Banana River, corresponding with the previously described dolphin community inhabiting portions of those two sub-basins [56]. A total of 55% of dolphins observed in the southern Indian River were only seen in this sub-basin, while the remaining animals were also seen in the northern Indian River and Banana River. The high proportion of individuals that exhibited site fidelity (55%) is consistent with dolphin communities previously described inhabiting portions of the southern Indian River [56], while movements into the Banana River and northern Indian River may account for seasonal variability observed in sub-basin abundance in the current study as well as in prior studies [19]. Future studies should investigate factors that influence dolphin movements in this sub-basin, including prey movement and environmental parameters that are known to contribute to dolphin movement in other regions [43–46].

Dolphin group size may be influenced by environmental factors, with shallow estuarine waters being predominated by smaller groups than open water habitats [62, 88–91]. Mean dolphin group size observed via aerial survey methods (2.45 ± 2.70 SD) [19] was somewhat smaller than was observed in the current study (4.08 ± 4.24 SD) which may be due to differences in survey methodology. Results were similar, however, to prior studies using similar platforms to evaluate IRL dolphin group size, which found an average group size of 4.1 ± 3.43 SD [92] and was further comparable to the mean group size observed in Sarasota Bay (4.8 ± 0.16 SE) [93]. The proportion of calves (including young of the year) observed during this study represented 19.2% of animals observed. While this proportion is significantly larger than prior abundance studies employed by line transect aerial survey which indicated calves represented 5.42% of the animals observed [19], differences are likely due to the conservative methods utilized to define a calf during aerial survey methods (half the size of the adult) [19]. Results are comparable to prior studies of Indian River Lagoon group composition which found that calves constituted 24% of individuals encountered [92]. Likewise, findings are similar to those observed in Sarasota Bay where calves constituted 21.5% of the population [94].

Unlike many previous studies of bay, sound and estuarine dolphins which occurred in more open settings [2, 13, 73, 86] the enclosed geography of the IRL [52] enabled survey effort of nearly the entire extent of available habitat within the study region, which prior studies have indicated is occupied by primarily residential dolphins [16–17]. Despite these extensive efforts, results from the current short-term capture-recapture study found an estimated 7% transient rate, using the criteria of a single sighting as an indicator. During the study, 84 marked dolphins (16.7%) were only sighted on one occasion and were considered to have very low site fidelity (potentially transient). These animals were distributed among sub-basins and seasons with 27% of cases occurring in Mosquito Lagoon. Examination of long-term photo-identification data revealed that the majority of dolphins that were only sighted once in Mosquito Lagoon (57%) were also known to inhabit the adjacent Halifax River estuary to the north and a few were known to range into the JES (S2 Table) [20]; therefore the complete range for these animals was not encompassed. However, the low transient rate suggested that transients do not play a large role in the population biology of IRL dolphins. The utilization of a parameter to account for transients did not influence abundance estimates (since these individuals were still included in calculations), however, the inclusion of this parameter resulted in more accurate survival rates for marked IRL adult residents by reducing the effect on apparent survival (reducing negative bias). It is possible that including the transient parameter may have biased resident survival high by removing individuals with the lowest survival from resident survival estimates [95]. This could be lessened in future studies by including covariates to identify animals known to range beyond the study area. The inclusion of a transient parameter, however, can aid in estimation in many cases and should be considered for similar estuarine Robust Design capture-recapture studies. In contrast to transience, temporary emigration occurred more frequently. The probability of an individual being absent from the study area during a primary period if it was absent in the session prior was greater (48%) than if it was present in the prior session (5%), suggesting that animals temporarily emigrated out of the study area and subsequently returned. Evidence for temporary emigration was mixed but generally supported non-structured movement (random or no movement) and provided evidence that some individuals were, at times, unavailable for recapture. Temporary emigration measured by the robust design can be due to either true absence from the study area or by animals being present but unobservable during a primary period. Because of the extensive sampling design and high detection within primary periods, it seems more likely that temporary emigration in this study was due to movement outside of the study area. Individuals inhabiting the edges of the study region with home ranges extending beyond the borders of the IRL may have contributed to temporary emigration during the study. While extensive efforts were undertaken to thoroughly cover the study area, individuals could have been unavailable in the labyrinth of canals and islands throughout the IRL. Additional causes for availability or perception bias [96] could be mitigated by including covariates influencing the detection process to reduce detection heterogeneity.

In three cases dolphins were thought to have low site fidelity (only sighted once and presumed transient), but were recovered dead and thus were true mortalities (S3 Table). During the study, estimated seasonal marked adult resident mortality rate ranged between 0.00-0.15, given a total of 503 marked adult dolphins this would be an estimated 0-75 marked adult mortalities. Documented marked adult mortality consisted of three dolphins (0.59%) that were sighted once and then unavailable for recapture (S3 Table). One of these animals (Hubbs-1654-Tt) was initially sighted in the southern Indian River near Sebastian Inlet (August 2016) and subsequently stranded alive on the Atlantic coast (October 2016), well north of the boundaries of the IRL (Ormond Beach), indicating the extensive range animals may traverse (S3 Table). It is likely that documented dolphin mortality is under-reported in the IRL and carcasses therefore unrecovered. A prior study indicated that only one third of dolphin carcasses were recovered in the well-studied Sarasota Bay estuary [97]. Utilizing this recovery rate to correct for undetected carcasses would adjust mortality to nine dolphin mortalities. The estimated mortality range (0-15%), while overlapping, estimates greatly exceeds the adjusted perceived mortality (1.8%). Estimated marked resident survival rates were broad across movement models, producing lower estimates that seemed unlikely for this population, as well as increased survival estimates that are more representative of documented survival for the Indian River Lagoon dolphin population. A trend of initially high resident survival, which decreased between the first (fall-winter) and the second (winter-spring) estimation interval was evident across all models that included both effects. While variation in survival between seasons might have occurred, the reduction in survival might instead be due to the limited number of primary periods in this study. Low-biased survival at the end of a study (the last survival estimates in a time series) has been noted in prior studies and diagnosed as the potential result of effects of constraints or improper estimates of parameters at the end of the time series [98]. Marked resident survival in the first time period (fall-winter), where it was likely modeled more precisely, ranged from 0.98-1.00 which more closely matched observed mortality. Future studies conducted over long timer periods with increased primary periods may be better suited to more accurately depict survival.

The mean proportion of marked dolphins (0.43 ± 0.12) was comparable but less than findings from other bay and estuarine systems that found a mean distinctiveness rate between 0.79 ± 0.09 and 0.72 ± 0.06 [13, 99] respectively. Differences in marked ratio between populations may be influenced by extrinsic and intrinsic factors related to differences in ecosystems types, as extensive portions of the Indian River Lagoon are relatively isolated from open water [52] compared to studies conducted in more open bays [2, 13, 73, 86]. Furthermore, in other study regions, rates of dorsal fin marking have been found to be influenced by sex, with males having significantly higher rates of dorsal fin nicks [100]. Potential heterogeneity in capture probability may occur as larger groups are more likely to contain calves [92] and by extension female animals, which may be less likely to be marked [100]. Sex-skewed marked ratios within the IRL population should be further examined. Interestingly, the marked ratio varied by sub-basin with the greatest ratio occurring in the southern Indian River (0.588) and the smallest ratio observed in the Banana River (0.294). Dolphins in the southern lagoon have the highest prevalence of boat-injuries, which results in dorsal fin disfigurement, disproportional to other sub-basins [101] and may account for some of the variability between sub-basins. On the contrary, much of the northern Banana River prohibits motorized vessels and is also further restricted to limited authorized personnel (no public usage) [102], thereby excluding anthropogenic activities (entanglement, vessel strikes and capture-release activities for dolphin health assessment) that may influence dorsal fin marking [8, 101, 103]. Detection probability ranged from 0.20 to 0.42 between secondary sessions which fell within recommended bounds of effective capture-recapture survey designs (0.2 - 0.3) [48]. Detection was the lowest in fall and was likely influenced by increased sea state, which is common in that season, causing perception bias [104]. Incorporating detection covariates for future efforts may aid to in improving precision and potentially reducing the required effort. Further investigation into factors that may influence dolphin detectability within the IRL including transience, ranging patterns beyond the study area, and potential perception and availability biases should be evaluated.

Effective management of the Indian River Lagoon Estuarine System dolphin stock requires current information on distribution and abundance. Estimates from this study provide the first comprehensive abundance estimates for IRL dolphins utilizing mark-recapture methodology and demonstrate the feasibility of utilizing robust design for this population. Likewise, the study provides the first estimates of temporary emigration, transient rate and survival. While the most reliable estimate of marked resident survival for Indian River Lagoon dolphins (0.98-1.00) during this short-term study was similar to survivorship estimates of bottlenose dolphins in other bays and estuaries (0.95 and 0.93) [29, 105], it should be taken into consideration that variability in survivorship may occur in this population as IRL dolphins have experienced reoccurring peaks in mortality (UMEs) (2001, 2008, 2013; 2013-2015) where annual mortality may range up to 77 dolphins (mean: 28.8 ± 2.48) [81]. Parameter estimates from this study will guide future survey design to provide estimates needed for stock assessment. Although conducted by different platforms, this study and prior aerial surveys revealed remarkably similar abundance estimates [19]. In recent years the occurrence of phytoplankton blooms and declining water clarity has become a growing concern in the IRL [36] and could influence dolphin availability for future estimates via line transect aerial survey. Therefore, future abundance estimates will need to rely on either ideal conditions for aerial survey methodology or the continued use of photo-identification mark-recapture methods utilizing Robust Design. Abundance estimates from the current study will enable stock managers to evaluate the impact of fisheries-related takes as well as the impact of mortality events. The data are timely as phytoplankton blooms and associated ecological impacts continue to threaten Indian River Lagoon health and the ramifications of this altered ecosystem are not yet known.

## Acknowledgements

We are very grateful to staff from the Kennedy Space Center Ecological Program Office, Kennedy Space Center, Cape Canaveral Air Force Station, and U.S. Fish and Wildlife Service for their assistance with entry into portions of the study area. The views expressed in this article do not necessarily represent the views of the National Aeronautics and Space Administration (NASA) or the United States. We thank Megan Stolen for assistance with stranding response and data collection. We sincerely thank our colleagues, Debbie Wingfield (Volusia County Environmental Management), Lisa Gemma, and St. Johns River Water Management District for providing critical field assistance and vessel support which made the project feasible. Likewise, we are grateful to those that were essential in the field as well as with photographic analyses: Amy Brossard, Heidy Clifford, Mackenzie Daniel, Agatha Fabry, Brandy Nelson, Samantha Nekolny, Jessie Stevens, and Elizabeth Titcomb. We sincerely appreciate Nicole Mader (Dolphin Ecology Project) enabling the study by providing essential coverage and photographic analyses in the southern end of the study area. Furthermore, we are grateful to the countless volunteers and interns who contributed to data collection and management. This study would not have been possible without our colleagues’ perseverance, talent and the dedication, for which we have the utmost gratitude. Lastly, we thank Lance Garrison for his assistance with the design of the project and continued guidance throughout the study.

## Supporting information

**S1 Text. Simulation study methods and results.**

**S1 Fig. Parameter estimates from Robust Design analysis of 1000 simulated data sets.** Dolphin populations were set with initial size 1000, a detection parameter of 0.3, survival at 0.95, and the rate of gamma prime (ϒ’= probability of a dolphin being unavailable for observation if unavailable in the prior primary period) and double gamma prime (ϒ’’= probability of a dolphin being unavailable if available in the prior primary period) were both set at 0.1. *Mean parameter estimate = dot, median = triangle, vertical line = generating parameter value.

**S1 Table. Estimated survival and movement parameters for the 12 best supported models for IRL bottlenose dolphins.** Estimates are calculated using profile likelihood estimation. To aid estimate comparisons, models with (a) and without (b) the transient parameter (time since first capture) are grouped. Time-specific adult survival rates (S) are presented for transients and residents, and survival estimates for time constant models are listed in the first time-specific row. Parameters include apparent survival (φ), model averaged estimates of adult survival rates (S) between primary periods, and temporary emigration: probability of an animal being temporarily unavailable for capture if the individual was available during the previous primary period (ϒ’’) or unavailable (ϒ’).

**S2 Table. Sighting history for thirteen bottlenose dolphins sighted only once in Mosquito Lagoon (considered potential transients).** Sighting data were collected between 2008 and the end of the current study. All animals listed had prior sightings north of the Indian River Lagoon in the Halifax River. *Indicate animals known to range north into the Jacksonville Estuarine System stock.

**S3 Table. Marked adult dolphins sighted during secondary sessions and subsequently recovered deceased during the study.**

